# Transient binding facilitates super-resolution imaging of functional amyloid fibrils on living bacteria

**DOI:** 10.1101/2025.06.19.660547

**Authors:** Daniel J. Foust, Divya Kolli, Kailyn Jessel, Zeyang Hu, Matthew R. Chapman, Julie S. Biteen

**Affiliations:** Department of Chemistry, University of Michigan, Ann Arbor, MI, United States; Department of Molecular, Cellular and Developmental Biology, University of Michigan, Ann Arbor, MI, United States

**Keywords:** CsgA, curli, *Escherichia coli*, biofilm, fluorescence microscopy, fluorescence correlation spectroscopy, super-resolution optical fluctuation imaging, single-molecule localization microscopy, Nile blue

## Abstract

Curli, which are the major proteinaceous component of the *Escherichia coli* biofilm extracellular matrix, help protect cells against environmental stressors including dehydration and antibiotics. Composed of the amyloid proteins CsgA and CsgB, curli self-assemble as these protomers are secreted into the extracellular space. The mechanisms of curli assembly and their functional roles within the extracellular matrix are incompletely understood. High-resolution imaging tools compatible with live-cell conditions provide a critical means to investigate the assembly and function of curli in their native context.

In this study, we use super-resolution imaging to visualize curli fibrils on living bacterial cells. Transient amyloid binding of the fluorogenic dye Nile blue facilitates two complementary super-resolution fluorescence microscopy approaches: single-molecule localization microscopy and super-resolution optical fluctuation imaging. Additionally, imaging fluorescence correlation spectroscopy was used to measure the characteristic relaxation times associated with Nile blue binding to CsgA fibrils. Together, these approaches offer a framework for imaging-based biophysical characterization of curli structures on living cells.

**Importance/impact statement:** *Escherichia coli* and other enteric bacteria secrete amyloid proteins that self-assemble into fibrillar structures called curli, forming a key component of biofilm extracellular matrices. Bacterial biofilms confer resilience to harsh environments with broad implications for human health. In this study, we extend transient amyloid binding to the novel application of super-resolution fluorescence microscopy of curli on living cells, offering promising approaches to gain structural and mechanistic insights.

## Introduction

*Escherichia coli* and other enteric bacteria secrete amyloid proteins that self-assemble into β-sheet-rich fibrils called curli (Chapman et al. 2002). Curli are the most abundant component of the *E. coli* extracellular matrix (ECM) that forms when bacterial communities form biofilms (McCrate et al. 2013). Bacterial biofilms contribute to disease pathogenesis and help confer resistance to environmental stressors including antibiotics (Kikuchi et al. 2005; Van Gerven et al. 2018). Additionally, curli have been linked to Parkinson’s disease pathology in mice, suggesting that they may play a role in potentiating neurodegenerative disease through interactions with host amyloids (Friedland and Chapman 2017; Sampson et al. 2020; Bhoite et al. 2022).

The amyloid protein CsgA is the major structural component of curli (Olsén et al. 1993). A second amyloid protein, CsgB, aids nucleation and anchoring of curli to the outer membrane (Bian and Normark 1997; Hammer et al. 2007). In addition to CsgA and CsgB, a set of accessory proteins regulate the transcription, transport, and conformation of the amyloids (Evans and Chapman 2014). The chaperone CsgC inhibits the amyloid fold and prevents intracellular fibrillization (Evans et al. 2015). CsgG forms an octameric pore to facilitate secretion across the outer membrane (Robinson et al. 2006). CsgE and CsgF support secretion through interactions with intracellular and extracellular surfaces of the pore, respectively (Nenninger et al. 2009; Klein et al. 2018; Swasthi et al. 2023). Curli production is tuned to a multitude of environmental factors including temperature, osmolarity, and nutrient availability (Andreasen et al. 2019). Understanding the structure and dynamics of curli fibers on living cells is crucial for elucidating their roles in both bacterial physiology and human health.

In this study, we extend advanced fluorescence microscopy techniques to the novel application of characterizing curli fibers on living cells. Here, fluorogenic dye Nile blue has been shown to transiently bind amyloid fibers *in vitro* (Ruiz-Arias et al. 2022). This behavior has been exploited to facilitate single-molecule localization microscopy (SMLM) in an approach known as transient amyloid binding (TAB) microscopy (Spehar et al. 2018; Sun et al. 2023). SMLM is based on the retrieval of the positions of single molecules from an image time series using algorithms to detect and localize single molecules (Tuson and Biteen 2015). Here, we have extended the TAB-SMLM approach for imaging intact curli assemblies on living cells.

SMLM encounters a limitation when the density of the target structure is highly variable, as is often the case *in cellulo*: under these conditions, it is challenging to tune the labeling strategy to effectively sample the entire structure. In a typical SMLM experiment, the TAB fluorogen concentrations need to be optimized to enable the detection of isolated fluorescent molecules in each imaging frame. However, in variable environments, TAB fluorogen concentrations that are calibrated to efficiently sample low-density regions, e.g., isolated fibrils, will result in high-density regions being oversampled such that single-molecule detection is rare. Conversely, if the fluorogen concentration is optimized for imaging high-density regions, the low-density regions will be under-sampled, requiring prohibitively long acquisition times to obtain an image that captures the full extent of the structure.

We therefore addressed these limitations of SMLM with TAB by implementing a second, complementary super-resolution approach: super-resolution optical fluctuation imaging (SOFI) (Dertinger et al. 2009). SOFI is based on the pixel-wise computation of cumulants to quantify correlations in the temporal fluctuations of the fluorescence signal (Dertinger et al. 2009). SOFI achieves increased image resolution based on the fact that, because the fluorescence from independent emitters is incoherent, their contributions to the final image are uncorrelated. SOFI does not rely on the detection and isolation of signals from single molecules, but it does require that the target switch between states with distinguishable fluorescence intensities (Dertinger et al. 2009). In the TAB experiments in this study, these distinguishable state changes are from the fluorogen binding and unbinding.

An additional benefit of SOFI is that imaging parameters can be chosen so that the data are also compatible with imaging fluorescence correlation spectroscopy (iFCS) (Kisley et al. 2015). iFCS is a camera-based analog of fluorescence correlation spectroscopy (FCS) that calculates autocorrelation functions for each pixel to make thousands of measurements in parallel (Cooper and Harris 2014). We implemented iFCS to measure characteristic fluorescence fluctuation relaxation times associated with Nile binding to curli on living cells. Our measurements support the presence of two characteristic fluctuation times for Nile blue binding, which may indicate multiple binding modes or multi-step binding as has been demonstrated for other dye/amyloid pairs with single-point measurements (Sarkar et al. 2023).

## Results

### Nile blue stains E. coli cells expressing curli

We sought a labeling strategy compatible with fluorescence imaging of curli fibers on living *E. coli* cells while leveraging the capabilities of super-resolution microscopy. We characterized Nile blue, a solvatochromic and fluorogenic dye that has been shown to transiently bind other amyloids, such as Aβ42 (Ruiz-Arias et al. 2022). Additionally, its excitation and emission in the red portion of the electromagnetic spectrum make it well-suited for low-background imaging *in cellulo* because the cellular autofluorescent background is higher in the blue region of the spectrum (Shcherbakova et al. 2012).

To test the affinity of Nile blue for the curli-producing *E. coli* strain MC4100, we supplemented agar plates with 100 µM Nile blue and cultured cells for 72 hours at room temperature. Under these conditions, wild-type cells formed biofilms rich in mature curli. These biofilms exhibited a blue hue, demonstrating their ability to accumulate Nile blue from the agar (Figure 1A). In contrast, a mutant strain lacking the amyloid protein genes *csgA* and *csgB* formed biofilms with a severely diminished capacity to retain the dye (Figure 1A). Notably, the presence of Nile blue in the agar did not prevent the growth of either strain, indicative of compatibility with live-cell assays.

**Figure 1.**
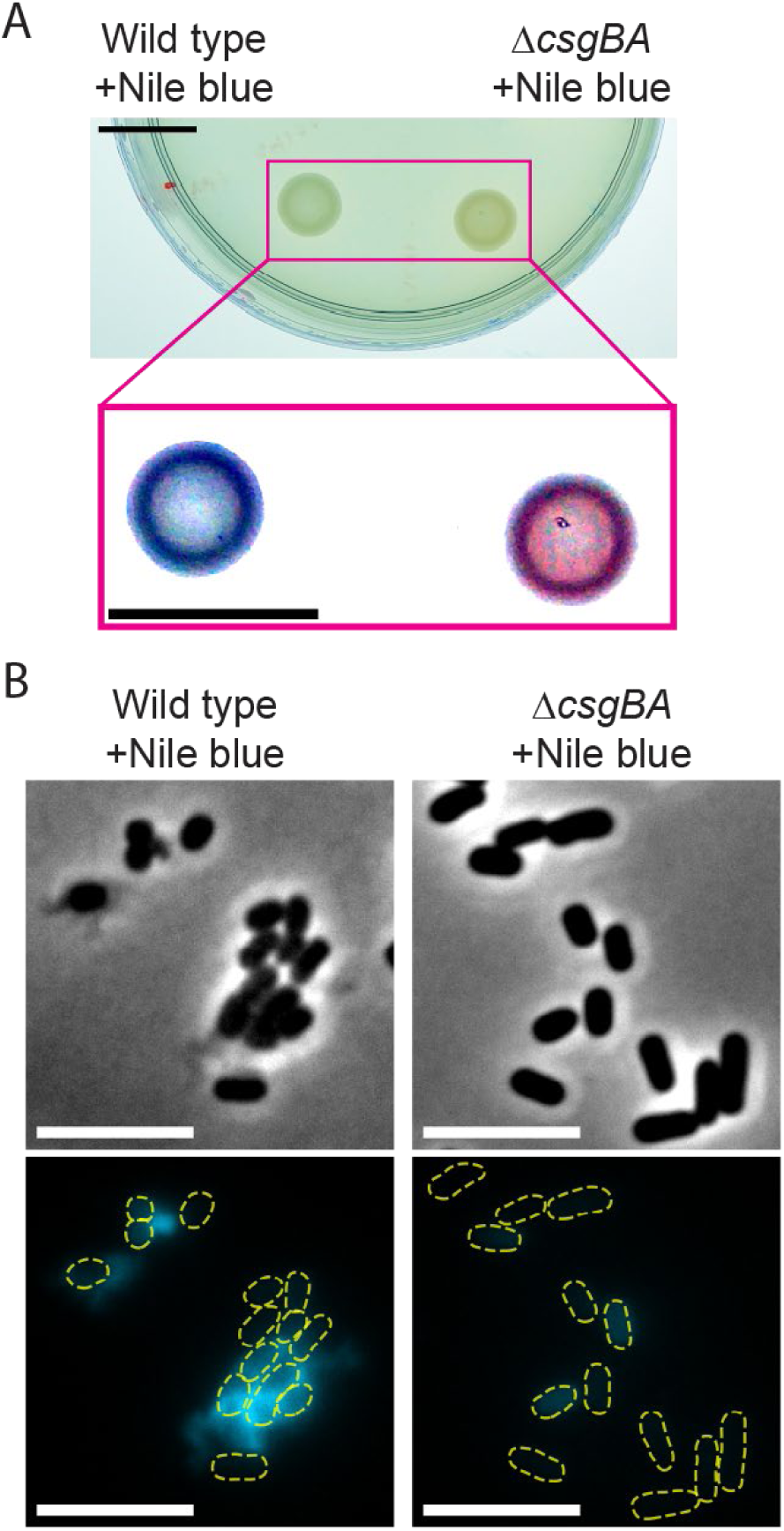
Nile blue stains *E. coli* cells that express curli. **(A)** *E. coli* biofilms grown on YESCA agar plates supplemented with Nile blue. The top image shows an unprocessed image captured with a smartphone camera. The outset below shows the same biofilms with background subtraction and the colors rebalanced to emphasize the difference in hue between the curli-expressing wild-type strain and the curli-deletion strain. Scale bars: 10 mm. **(B)** *E. coli* cells dispersed from biofilms. Top row: phase-contrast images. Bottom row: Nile blue fluorescence. Yellow dashed lines indicate cell outlines segmented from the phase-contrast images above. Scale bars: 5 µm.

We visualized cells that had been grown on agar plates without Nile blue using epifluorescence microscopy. We added Nile blue to the resuspended cells just before pipetting them onto agarose pads for imaging. Under 640-nm laser excitation, wild-type *E. coli* exhibited robust Nile blue fluorescence that extended away from the cell bodies, consistent with the labeling of extracellular structures (Figure 1B). Nile blue fluorescence was associated with dimmer cell-adjacent regions in the phase-contrast images, consistent with the presence of optically dense extracellular structures such as curli (Figure 1B). In the Δ*csgAB* strain, we observed only low-intensity Nile blue fluorescence originating from cell bodies, which we attribute to low-affinity, non-specific binding of Nile blue to the cell surface (Figure 1B), indicating that the significant off-cell fluorescence in the wild-type cells is indeed attributable to curli amyloid labeling.

### Transient binding of Nile blue enables live-cell SMLM

We investigated the potential for transient binding of Nile blue to facilitate super-resolution fluorescence microscopy of curli on living *E. coli* (Figure 2A). We considered two super-resolution techniques, illustrated conceptually using simulated data in Figure 2: single-molecule localization microscopy (SMLM) (Figure 2B-C) and super-resolution optical fluctuation imaging (SOFI) (Figure 2D-F). SMLM is enabled by the detection and sub-pixel localization of single molecules (Figure 2B-C) while SOFI relies on the statistical analysis of fluorescence fluctuations (Figure 2D-F). Fluorescence fluctuations generated from Nile blue brightening upon binding to curli can be leveraged to support both approaches.

**Figure 2.**
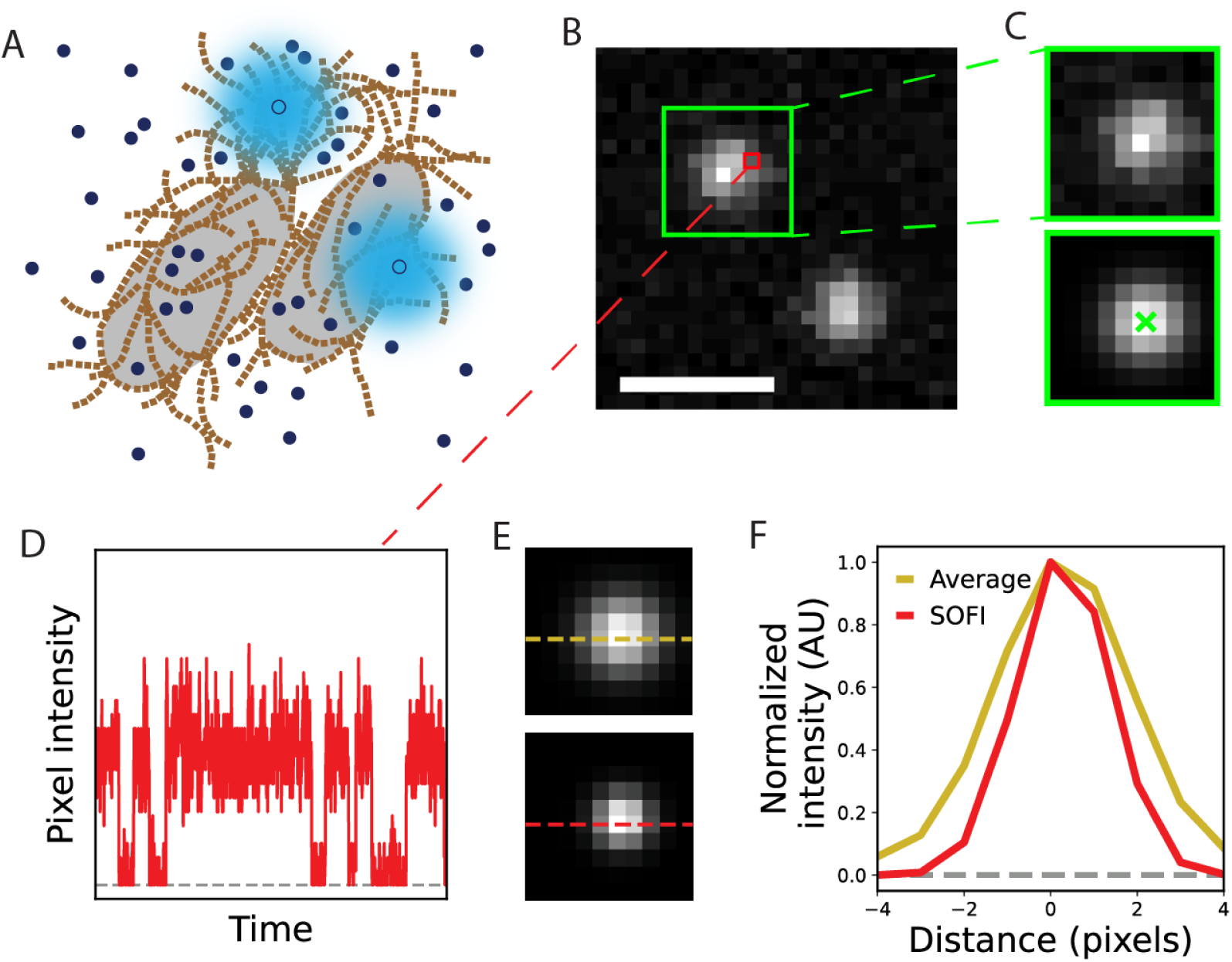
Transient amyloid binding enables super-resolution microscopy on living *E. coli*. **(A)** Schematic representation of two *E. coli* cells (light grey) expressing extracellular curli (light brown). Unbound Nile blue molecules (dark blue) remain in a dark state, while curli-bound Nile blue molecules (light blue) fluoresce brightly. **(B)** Simulated image of fluorescence from two fluorescent molecules (green box). Scale bar: 1 µm. **(C)** SMLM is based on the detection and subpixel localization of single molecules. Top: sub-images excised from a single frame (B, green box) are fit with a model PSF (bottom) to determine the molecule position (green ‘×’) with sub-diffraction-limit precision. **(D)** SOFI is based on temporal fluorescence intensity fluctuations (changes in brightness) analyzed for each pixel such as the one outlined in red in B. **(E)** Temporal average (top) and SOFI reconstruction (bottom) for the molecule in the green box in B over many imaging frames. **(F)** Normalized intensity profiles along the dashed lines in E show the localization precision enhancement from SOFI.

SMLM requires sparse labeling so that the diffraction-limited emission patterns of nonoverlapping emitters can be temporally isolated when imaged on a camera and precisely fit with a model point-spread function (PSF), commonly a 2D Gaussian distribution (Figure 2A-C). This requirement can be satisfied with transient amyloid binding by tuning the concentration of the probe dye so that only a few dye molecules are bound to the structure of interest per imaging frame (Figure 2A). Then, over thousands of imaging frames, many binding events are detected, super-localized, and used to construct a super-resolution density map (Tuson and Biteen 2015).

SMLM of curli extending from cells dispersed from an *E. coli* biofilm is demonstrated in Figure 3. Cells were identified using phase-contrast microscopy (Figure 3A) and imaged with 30-ms imaging frame times (Movie S1). For comparison, we computed a diffraction-limited image from the temporal average of all frames in the movie (Figure 3B). The SMLM image reveals fine heterogeneities along the cell surface and in the expansive extracellular curli mass that are not apparent in the diffraction-limited image (Figure 3C).

**Figure 3.**
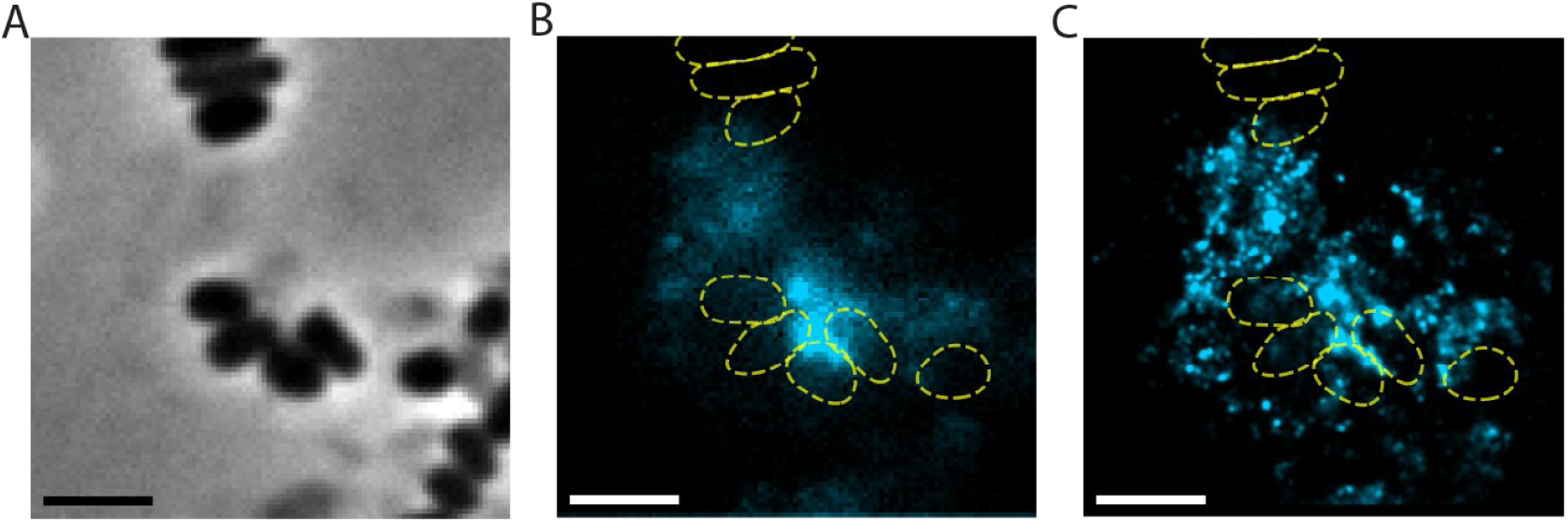
Single-molecule localization microscopy via transient binding of Nile blue. **(A)** Phase-contrast image of dispersed cells expressing curli. **(B)** Temporal average of Nile blue fluorescence. Colorscale: fluorescence intensity. **(C)** Super-resolution density map of Nile blue binding using SMLM. Colorscale: localization density. Yellow dashed lines are outlines of cells found from the segmentation of cells in the phase-contrast image in A. Scale bars: 2 µm.

### SOFI and iFCS of Nile blue binding to curli on live E. coli

SOFI is based on a pixelwise temporal correlation analysis of raw image time series (Figure 2B-C). The technique relies on switching between brightness states of the target (Figure 2D). In these transient curli binding experiments, the target curli fibers are nonfluorescent under 640-nm excitation (low brightness), and fluorescence is detected when Nile blue binds (high brightness) (Figure 2A, D). The fluorescence emission of unique emitters is uncorrelated due to the independence of the Nile blue binding and unbinding events. Pixelwise autocorrelation is quantified through the calculation of cumulants, filtering out contributions from uncorrelated signals. We calculated second-order cumulants resulting in a squaring of the effective point spread function and a 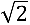 improvement in the resolution in the SOFI image in comparison to its diffraction-limited counterpart (Figure 2E-F) (Dertinger et al. 2009).

Cells expressing curli from dispersed *E. coli* biofilms were identified using phase-contrast microscopy (Figure 4A). We measured temporal fluorescence fluctuations from Nile blue binding to curli fibers on living *E. coli* cells by collecting movies with short (2.5-ms) exposure times (Movie S2). Although the simulated data in Figure 2D represents the binding and unbinding of exactly one molecule at a time for simplicity, SOFI does not require that fluorescence fluctuations be generated from single molecules. In fact, SOFI is generally applicable over a broad range of dye concentrations and binding densities (Movie S2). We used these movies to generate super-resolved images of curli fibers via a SOFI reconstruction algorithm (Dertinger et al. 2009). In contrast to the diffraction-limited temporal average (Figure 4B), the SOFI map (Figure 4C) reveals the dense, irregular extracellular meshwork formed by curli with much greater detail.

**Figure 4.**
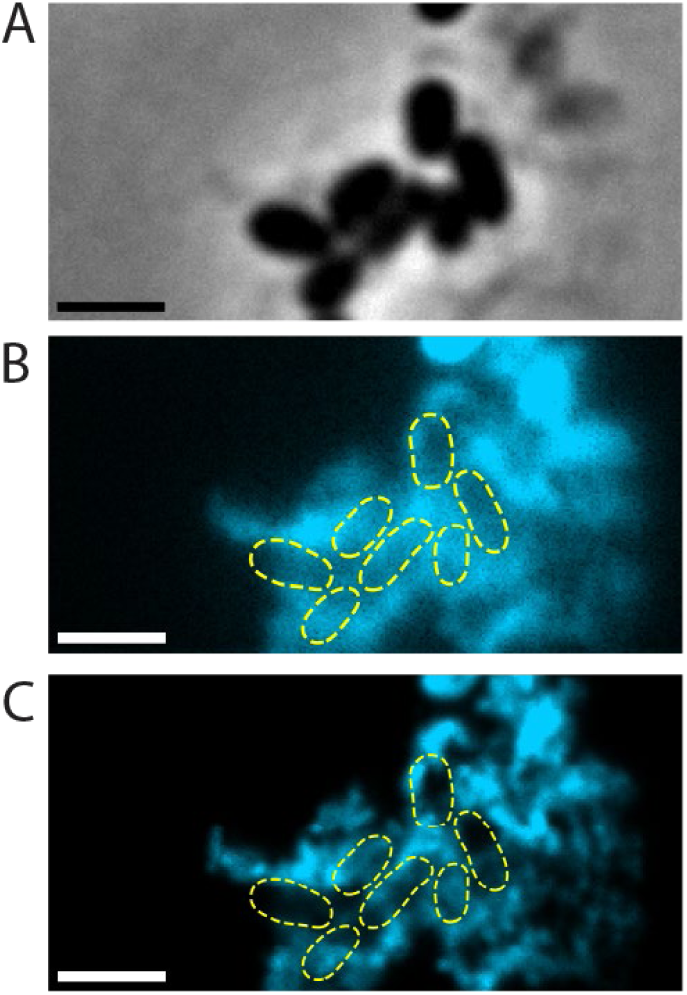
Super-resolution optical fluctuation imaging via transient binding of Nile blue. **(A)** Phase-contrast image of dispersed *E. coli*. **(B)** Temporal average of Nile blue fluorescence. Colorscale: fluorescence intensity. **(C)** SOFI reconstruction. Colorscale: SOFI density. Dashed yellow outlines indicate cell positions determined by segmentation of the phase-contrast image in A. Scale bars: 2 µm.

Furthermore, the fluorescence intensity traces provide information to characterize the relaxation times of Nile blue on curli using iFCS. To perform iFCS, we calculated the autocorrelation function from the temporal fluorescence intensity traces from each pixel (Figure 5A). We fit individual autocorrelation functions with model functions using least squares optimization (Figure 5B). Generally, a single-component model was inadequate for fitting the autocorrelation functions, as evaluated using reduced-*χ*^2^ statistics. To better capture the observed dynamics, we used a two-component model, yielding average fast and slow relaxation times of 23 ± 1 ms (mean ± SEM) and 482 ± 13 ms, respectively. The relaxation time distributions are shown in Figure 5C.

**Figure 5.**
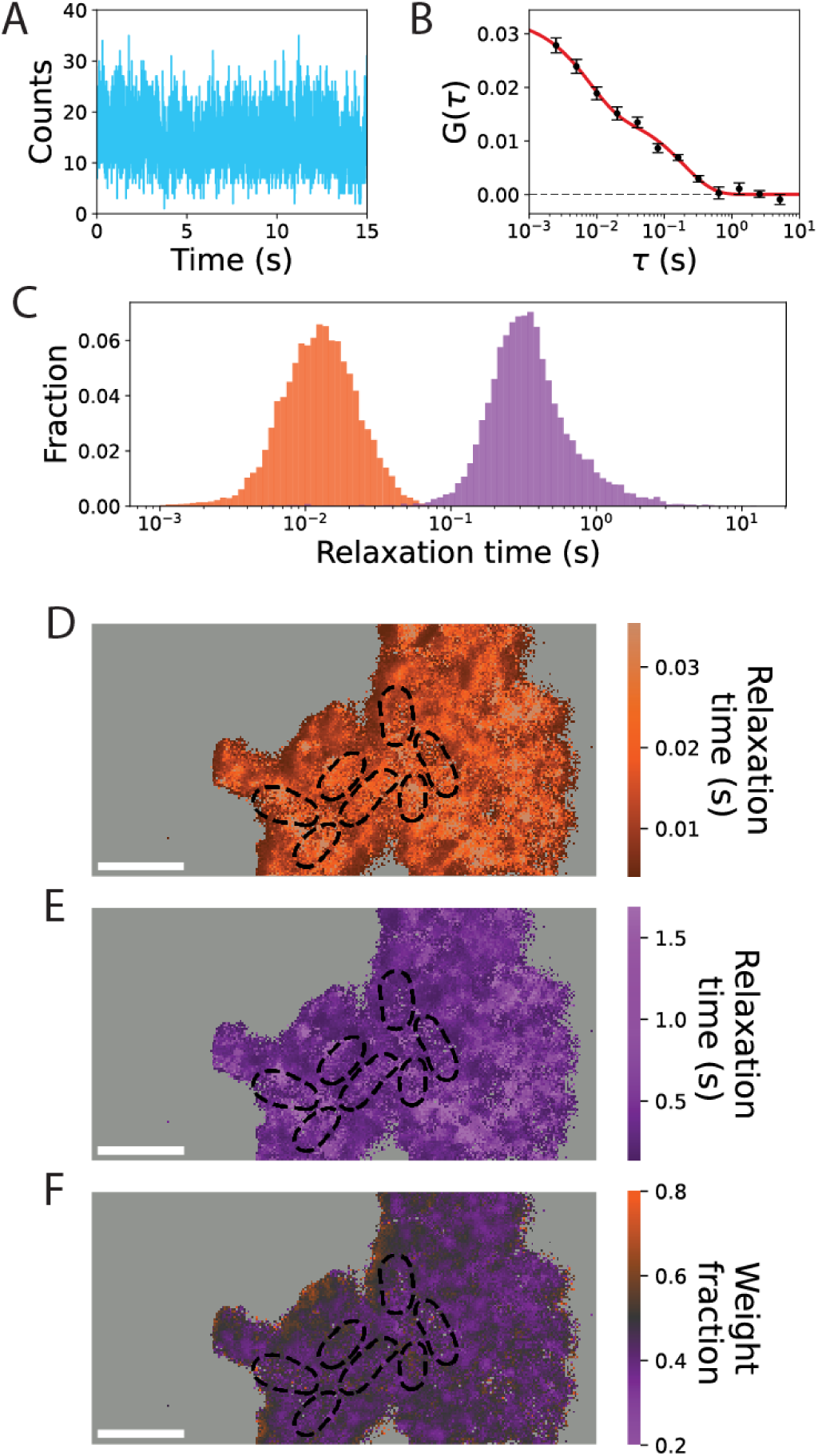
Imaging fluorescence correlation spectroscopy measures Nile blue binding kinetics. **(A)** Fluorescence intensity trace for a representative pixel from the same movie used to generate the SOFI image in Figure 4. **(B)** Average autocorrelation function for the same pixel over six 15-s movie segments calculated as a function of timelapse, *τ*. Black circles indicate the mean of the autocorrelation functions for that pixel and error bars indicate the standard error of the mean. The red line is a fit to a two-component binding model. **(C)** Histogram of characteristic relaxation times, determined by fitting the autocorrelation functions for each pixel with SNR > 10. Shorter and longer relaxation times are shown in orange and purple, respectively. **(D) – (F)** Maps of parameters obtained from fitting the autocorrelation function for each pixel. **(D)** Map of short relaxation times. **(E)** Map of long relaxation times. **(F)** Map of the estimated fraction of molecules associated with the shorter relaxation time. Black dashed lines indicate segmented cell outlines. Pixels with insufficient SNR for correlation analysis are colored light grey. Scale bars: 2 µm.

We use the general term relaxation time because the sources of the intensity fluctuations cannot be identified unambiguously. Previously, the presence of two components in point FCS measurements on fluorogenic dyes binding to amyloids has been used as evidence of two residence times, each associated with a distinct binding mode (Sarkar et al. 2023). Multimodal binding may also be characteristic of Nile blue binding to CsgA fibrils, however, we cannot exclude the possibility that shorter relaxation time corresponds to blinking associated with a photophysical phenomenon (Schenk et al. 2004; M et al. 2023).

iFCS offers advantages over classic single-point FCS measurements by enabling thousands of measurements to be made in parallel and, consequently, by spatially resolving these measurements. Leveraging these advantages, we generated maps of characteristic binding times (Figure 5D-E) and their associated weight fractions (Figure 5F). Weight fractions estimate the fraction of bound molecules associated with each relaxation time.

The relaxation times and their associated weight fractions did not generally follow any clear spatial organization relative to the positions of cells (Figure 5D-F), though we observed that both the fast and slow relaxation times are lower (Figure 5D-E) in the regions of dimmer fluorescence intensity (Figure 4B-C) that tend to be at the edges of the region of interest. This correspondence is most likely an artifact: autocorrelation function fitting with least-squares optimization is known to be a biased estimator that underestimates relaxation times in photon-poor regions (Kohler et al. 2023). Accordingly, the weight fraction associated with the shorter relaxation time is higher in these regions (Figure 5F).

## Discussion

We have demonstrated that the fluorogenic dye Nile blue is readily absorbed by *E. coli* expressing extracellular amyloid fibrils known as curli (Figure 1). Furthermore, we showed that transient binding of Nile blue is an effective tool for imaging curli on living cells (Figures 2 – 4). A major advantage of TAB relative to other labeling strategies is that it does not require covalent modification of the protein of interest. This feature is especially important in the context of amyloids, as covalent modifications may alter their aggregation kinetics (Bhoite et al. 2023). We demonstrated that this flexible, non-covalent labeling approach can be leveraged for both SMLM (Figure 3) and SOFI (Figure 4). Additionally, by coupling SOFI with iFCS, we measured autocorrelation functions that suggest two distinct relaxation times associated with Nile blue binding to CsgA fibrils (Figure 5).

SMLM and SOFI each have strengths and weaknesses. SMLM is the higher resolution approach, as the precision with which single-molecule positions can be determined is primarily limited by the number of detected photons (Thompson et al. 2002). Typical SMLM resolutions are 10 – 30 nm (Sage et al. 2019). In SOFI, the achievable enhancements in resolution are proportional to the square root of the cumulant order (Dertinger et al. 2009). In this work, we used second-order cumulants, yielding approximately a 1.4-fold resolution enhancement (to ∼150 nm). Although higher-order cumulants can provide greater resolution, they require more data and are more susceptible to artifacts (Basak et al. 2025), though a recent advance leverages an artificial neural network with a transformer architecture to achieve SMLM-like resolution using SOFI-like inference from temporal correlations (Reinhard et al. 2025).

Another consideration in super-resolution microscopy is the set of inherent tradeoffs in acquisition time, temporal resolution, and spatial sampling. The conventional SMLM detection and localization scheme used in this work requires that the emission patterns of single molecules be well-separated on the camera for accurate determination of molecular positions. Although we were able to detect single molecules in both the curli-dense interstitial spaces between cells and the sparser regions away from cells (Figure 3C, Movie S1), navigating this dynamic range required us to choose a 1 µM Nile blue concentration that was not optimal for either extreme. In the most curli-dense regions, the dye concentration was higher than optimal, and overlapping Nile blue binding events led to errors in single-molecule detection and localization (Movie S1). Conversely, the relatively low dye concentration required longer acquisition times to adequately sample the more sparse regions. Sophisticated algorithms that support multi-emitter fitting (Holden et al. 2011; Huang et al. 2011) or leverage neural networks trained to extract localizations from overlapping emitters (Nehme et al. 2018; Speiser et al. 2021; Reinhard et al. 2025) can partially alleviate these limitations in SMLM. In contrast, SOFI does not inherently necessitate limitations on the emitter density. Reconstructions have been achieved using as few as 20 frames, and continued improvements in SOFI algorithms remain an area of active research (Zhao et al. 2023; Basak et al. 2025).

In the context of imaging bacterial amyloids, the choice between SMLM and SOFI depends on the desired biological information. SOFI is strongest when high temporal resolution or high throughput is desired and when the additional resolution enhancement afforded by SMLM is not necessary. In addition to providing better resolution, SMLM can be extended with polarization resolution to measure single-molecule orientations, a technique known as single-molecule orientation localization (Backlund et al. 2014). Single-molecule orientations provide readouts of *in situ* structural biology. The orientation of Nile blue has been used to probe the amyloid Aβ42 (Sun et al. 2023).

To our knowledge, this is the first application of iFCS to spatially resolve transient amyloid binding kinetics. CsgA fibrils are known to take on different conformations in response to environmental conditions (Andreasen et al. 2019; Bu et al. 2024). We posit that iFCS in conjunction with amyloid-sensitive fluorogens whose binding kinetics are sensitive to fibril structure could be used to probe structural variations in intact biofilms.

Recently, covalent labeling of CsgA with fluorescent tags has been used to quantify elongation rates *in vitro* (Olsen et al. 2024; Olsen, Christensen, et al. 2025; Olsen, Larsen, et al. 2025). It is not known whether these findings translate to living cells, where the presence of CsgB— the nucleation-inducing curli protein—and the complexity of the cellular milieu may significantly alter assembly dynamics. The imaging approaches explored in this work present opportunities to probe curli assembly *in vivo*.

## Materials and methods

### Cell preparation

We cultured *E. coli* strain MC4100 by streaking onto agar plates containing yeast extract and casamino acids (YESCA), supplemented with 100 µM Nile blue where indicated. We used a mutant strain with the *csgBA* operon deleted (LSR13) as a negative control. We grew cultures for 48 – 72 h at room temperature. We recorded macroscopic images of agar plates by illuminating from below with a white light source and capturing photographs with a Samsung S22 smartphone. Images were processed using ImageJ. To control the initial cell density in plate imaging experiments, a 4 µL spot of overnight culture diluted to an optical density of 1.0 was pipetted directly onto the agar plate for each strain.

We observed that cells grown on Nile blue-supplemented plates exhibited a higher degree of nonspecific fluorescence under 640-nm excitation, therefore, for imaging dispersed cells, we grew cells on plates without Nile blue and added Nile blue just prior to preparation for imaging. To prepare cells for microscopy, we scooped a lentil-sized amount of the biofilm using a disposable sterile loop and transferred it to 500 µL of phosphate buffered saline (PBS). We pipetted vigorously to disperse the clump into single cells and small cell clusters. We added Nile blue to a concentration of 1 µM or 5 µM for SMLM and SOFI experiments, respectively. 2 – 5 µL of cells were pipetted onto a 2% w/v agarose pad. We covered the agarose pad with a #1.5H thickness cover glass (Thorlabs) for microscopy.

### Fluorescence microscopy

We imaged live *E. coli* samples on a custom-built optical instrument configured for epifluorescence microscopy. To excite Nile blue, we used 640-nm continuous-wave laser excitation (Coherent CUBE 640-40C) cleaned up with an excitation filter (Chroma, Z640). We directed the beam through the back aperture of an Olympus IX71 microscope and focused it on the back focal plane of a 100×, 1.45 numerical aperture objective lens (Olympus X-Apo). We used a dichroic mirror (Chroma Di01-R640) to direct collimated light onto the sample. We used a combination of linear polarizer (Thorlabs LPVISB-050-M), half-wave plate (Thorlabs AHWP10M-600), and liquid crystal variable retarder (Thorlabs LCC1223T-A) to correct for the phase shift introduced by the dichroic mirror to produce circularly polarized light at the sample. The average power density within the full width at half maximum diameter of the Gaussian beam at the sample plane was approximately 10 µW/µm^2^. We collected emitted fluorescence with the same objective lens, filtered it by transmission through the dichroic mirror and a long pass filter (Semrock, BLP01-635R-25), and directed it onto an ORCA-Quest qCMOS camera (Hamamatsu C15550-20UP) positioned at the image plane of the IX71 tube lens. The pixel size at the object plane was 46 nm, except when 2×2 binning was used for SMLM. We operated the camera in ultra quiet mode with photon number resolving for low-intensity imaging (less than 200 photons/pixel/frame) and switched off photon number resolving for higher-intensity imaging due to the limited dynamic range.

### Single-molecule localization microscopy (SMLM)

We recorded movies with 30-ms exposure time for 2000-10000 frames per movie. Typically, multiple movies were collected per field-of-view. 2×2 pixel binning was used so that the pixel size at the object plane was 92 nm.

We detected and localized single molecules using Palmari, a plugin for the Python-based image analysis software Napari (Sofroniew et al. 2025). To detect single molecules, we applied a Laplacian-of-Gaussian filter (*σ* = 1.5 pixels) to raw movie frames (Godinez et al. 2009). We identified candidate molecules by applying a threshold to each filtered frame equal to three times the standard deviation of the filtered image pixel values. We fit sub-images of single molecules with a 2D-Gaussian PSF using GPUfit (Przybylski et al. 2017). We discarded candidate molecules if a high-quality fit could not be achieved (Isaacoff et al. 2019).

We constructed super-resolution density maps by generating 2D histograms of all localizations with 5-nm × 5-nm bins. We convolved these histograms with a 2D Gaussian kernel with *σ* = 30 nm selected to correspond to the estimated average localization uncertainty to produce the final image.

### Super-resolution optical fluctuation imaging (SOFI)

For each field of view, we recorded movies with 2.5-ms exposure times with a qCMOS camera operating in photon number resolving mode. The principle of SOFI is to enhance spatial resolution by applying temporal correlation analysis to raw image time series. This approach leverages the fact that the fluorescence emission of unique emitters is uncorrelated. By performing pixel-wise calculations of cumulants, individual emitter contributions are independent in the final image.

In this work, we used second-order cumulants to generate SOFI images:

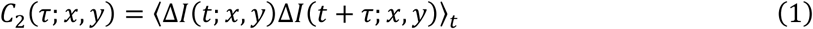

Where: *ΔI*(*t*) = *I*(*t*) − 〈*I*(*t*)〉_*t*_. We summed second-order cumulants for many time lags, *τ*, to generate SOFI images, *I*_*x*,*y*_ *^SOFI^*:

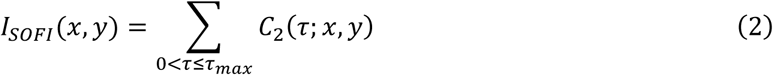

We chose *τ_max_* to be the point at which the autocorrelation function had decayed by ∼50% to avoid noise contributions from larger time lags. Movies were divided into 15-s segments and we averaged SOFI images calculated for each segment to produce the final image.

### Imaging fluorescence correlation spectroscopy (iFCS)

Using the same movies used for generating SOFI images, we calculated the temporal autocorrelation function, *G*(*τ*; *x*, *y*), from the intensity trace, *I*(*t*; *x*, *y*), for each pixel:

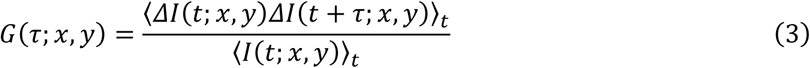

Notably, the numerator in Equation (3) is equivalent to Equation (1), allowing computations to be shared between SOFI and iFCS when applied to the same data.

We averaged autocorrelation curves from three to six 15-s movie segments and used the standard error of the mean (SEM) to quantify the uncertainty at each time lag.

We fit experimental autocorrelation functions, *G*(*τ*), with a two-component binding model:

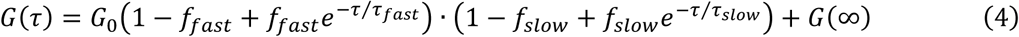

*G*_0_ is the amplitude and is inversely proportional to the average number of molecules. *τ_fast_* and *τ_slow_* are binding relaxation times and *f_fast_* and *f_slow_* are the corresponding fractions of bound molecules associated with each relaxation time. *G*(∞) is the offset.

We also considered a single-component model that can be derived from Equation (4) by setting *f_fast_* = 1 and *f_slow_* = 0. We evaluated models by calculating reduced-*χ*^2^ statistics from residuals of the least squares optimization. Generally, we calculated reduced-*χ*^2^ ≫ 1 from fitting the single-component model, indicating an inadequate description of the data, and reduced-*χ*^2^ ≈ 1 for the two-component model.

To generate maps of the fit parameters, results from low signal-to-noise (SNR) autocorrelation functions were excluded. SNR was estimated by dividing the autocorrelation function amplitude by the standard deviation of the last five time lags of the autocorrelation function. Results from autocorrelation functions with SNRs below 10 are not shown on maps and are substituted with light grey pixels.

## Supplementary material description

**Movie S1: Single-molecule localization microscopy via transient binding of Nile blue.** A subset of frames from the movie used to create Figure 3B-C. The 33-fps playback speed is equal to the acquisition speed. Scale bar: 2 µm.

**Movie S2: Super-resolution optical fluctuation imaging via transient binding of Nile blue.** A subset of frames from the movie used to create Figures 4-5. The 40-fps playback speed is one-tenth of the 400-fps acquisition speed. Scale bar: 2 µm.

## Acknowledgments

This work was supported by NSF Grant CHE-2403937 to JSB and NIH Grant R01 GM118651 to MRC. We thank Dr. Matthew Lew for consultation on the selection of Nile blue for these experiments, Dr. Saaj Chattopadhyay for supporting the maintenance of the optical instrumentation, Adam Decker and Dr. Zechariah Pfaffenberger for experimental perspective from their unpublished pilot studies on imaging curli with SMLM, and Sarah El-Mohri for providing technical support.

## Abbreviations and symbols

E. coli: Escherichia coli
ECM: extracellular matrix
FCS: fluorescence correlation spectroscopy
iFCS: imaging FCS
PBS: phosphate buffered saline
PSF: point-spread function
SMLM: single-molecule localization microscopy
SNR: signal-to-noise ratio
SOFI: super-resolution optical fluctuation imaging
TAB: transient amyloid binding
YESCA: yeast extract supplemented with casamino acids

## Notes

### Competing Interest Statement

The authors have declared no competing interest.

## References

Andreasen M, Meisl G, Taylor JD, Michaels TCT, Levin A, Otzen DE, Chapman MR, Dobson CM, Matthews SJ, Knowles TPJ. 2019. Physical Determinants of Amyloid Assembly in Biofilm Formation. mBio. 10(1):10.1128/mbio.02279-18. doi:10.1128/mbio.02279-18.

Backlund MP, Lew MD, Backer AS, Sahl SJ, Moerner WE. 2014. The Role of Molecular Dipole Orientation in Single-Molecule Fluorescence Microscopy and Implications for Super-Resolution Imaging. ChemPhysChem. 15(4):587–599. doi:10.1002/cphc.201300880.

Basak S, Chizhik A, Gallea JI, Gligonov I, Gregor I, Nevskyi O, Radmacher N, Tsukanov R, Enderlein J. 2025. Super-resolution optical fluctuation imaging. Nat Photon. 19(3):229–237. doi:10.1038/s41566-024-01571-3.

Bhoite SS, Han Y, Ruotolo BT, Chapman MR. 2022. Mechanistic insights into accelerated α-synuclein aggregation mediated by human microbiome-associated functional amyloids. Journal of Biological Chemistry. 298(7). doi:10.1016/j.jbc.2022.102088. https://www.jbc.org/article/S0021-9258(22)00529-4/abstract.

Bhoite SS, Kolli D, Gomulinski MA, Chapman MR. 2023. Electrostatic interactions mediate the nucleation and growth of a bacterial functional amyloid. Front Mol Biosci. 10. doi:10.3389/fmolb.2023.1070521. https://www.frontiersin.org/journals/molecular-biosciences/articles/10.3389/fmolb.2023.1070521/full.

Bian Z, Normark S. 1997. Nucleator function of CsgB for the assembly of adhesive surface organelles in Escherichia coli. The EMBO Journal. 16(19):5827–5836. doi:10.1093/emboj/16.19.5827.

Bu F, Dee DR, Liu B. 2024. Structural insight into Escherichia coli CsgA amyloid fibril assembly. mBio. doi:10.1128/mbio.00419-24. https://journals.asm.org/doi/10.1128/mbio.00419-24.

Chapman MR, Robinson LS, Pinkner JS, Roth R, Heuser J, Hammar M, Normark S, Hultgren SJ. 2002. Role of Escherichia coli Curli Operons in Directing Amyloid Fiber Formation. Science. 295(5556):851–855. doi:10.1126/science.1067484.

Cooper JT, Harris JM. 2014. Imaging Fluorescence-Correlation Spectroscopy for Measuring Fast Surface Diffusion at Liquid/Solid Interfaces. Anal Chem. 86(15):7618–7626. doi:10.1021/ac5014354.

Dertinger T, Colyer R, Iyer G, Weiss S, Enderlein J. 2009. Fast, background-free, 3D super-resolution optical fluctuation imaging (SOFI). PNAS. 106(52):22287–22292. doi:10.1073/pnas.0907866106.

Evans ML, Chapman MR. 2014. Curli biogenesis: Order out of disorder. Biochimica et Biophysica Acta (BBA) -Molecular Cell Research. 1843(8):1551–1558. doi:10.1016/j.bbamcr.2013.09.010.

Evans ML, Chorell E, Taylor JD, Åden J, Götheson A, Li F, Koch M, Sefer L, Matthews SJ, Wittung-Stafshede P, et al. 2015. The Bacterial Curli System Possesses a Potent and Selective Inhibitor of Amyloid Formation. Molecular Cell. 57(3):445–455. doi:10.1016/j.molcel.2014.12.025.

Friedland RP, Chapman MR. 2017. The role of microbial amyloid in neurodegeneration. PLOS Pathogens. 13(12):e1006654. doi:10.1371/journal.ppat.1006654.

Godinez WJ, Lampe M, Wörz S, Müller B, Eils R, Rohr K. 2009. Deterministic and probabilistic approaches for tracking virus particles in time-lapse fluorescence microscopy image sequences. Medical Image Analysis. 13(2):325–342. doi:10.1016/j.media.2008.12.004.

Hammer ND, Schmidt JC, Chapman MR. 2007. The curli nucleator protein, CsgB, contains an amyloidogenic domain that directs CsgA polymerization. Proceedings of the National Academy of Sciences. 104(30):12494–12499. doi:10.1073/pnas.0703310104.

Holden SJ, Uphoff S, Kapanidis AN. 2011. DAOSTORM: an algorithm for high-density super-resolution microscopy. Nat Methods. 8(4):279–280. doi:10.1038/nmeth0411-279.

Huang F, Schwartz SL, Byars JM, Lidke KA. 2011. Simultaneous multiple-emitter fitting for single molecule super-resolution imaging. Biomed Opt Express, BOE. 2(5):1377–1393. doi:10.1364/BOE.2.001377.

Isaacoff BP, Li Y, Lee SA, Biteen JS. 2019. SMALL-LABS: Measuring Single-Molecule Intensity and Position in Obscuring Backgrounds. Biophysical Journal. 116(6):975–982. doi:10.1016/j.bpj.2019.02.006.

Kikuchi T, Mizunoe Y, Takade A, Naito S, Yoshida S. 2005. Curli Fibers Are Required for Development of Biofilm Architecture in Escherichia coli K-12 and Enhance Bacterial Adherence to Human Uroepithelial Cells. Microbiology and Immunology. 49(9):875–884. doi:10.1111/j.1348-0421.2005.tb03678.x.

Kisley L, Brunetti R, Tauzin LJ, Shuang B, Yi X, Kirkeminde AW, Higgins DA, Weiss S, Landes CF. 2015. Characterization of Porous Materials by Fluorescence Correlation Spectroscopy Super-resolution Optical Fluctuation Imaging. ACS Nano. 9(9):9158–9166. doi:10.1021/acsnano.5b03430.

Klein RD, Shu Q, Cusumano ZT, Nagamatsu K, Gualberto NC, Lynch AJL, Wu C, Wang W, Jain N, Pinkner JS, et al. 2018. Structure-Function Analysis of the Curli Accessory Protein CsgE Defines Surfaces Essential for Coordinating Amyloid Fiber Formation. mBio. 9(4):10.1128/mbio.01349-18. doi:10.1128/mbio.01349-18.

Kohler J, Hur K-H, Mueller JD. 2023. Autocorrelation function of finite-length data in fluorescence correlation spectroscopy. Biophysical Journal. 122(1):241–253. doi:10.1016/j.bpj.2022.10.027.

M M, Patidar RK, Tiwari R, Srivastava N, Ranjan N. 2023. Nile Blue: A Red-Emissive Fluorescent Dye That Displays Differential Self-Assembly and Binding to G-Quadruplexes. J Phys Chem B. 127(46):9915–9925. doi:10.1021/acs.jpcb.3c05084.

McCrate OA, Zhou X, Reichhardt C, Cegelski L. 2013. Sum of the Parts: Composition and Architecture of the Bacterial Extracellular Matrix. Journal of Molecular Biology. 425(22):4286–4294. doi:10.1016/j.jmb.2013.06.022.

Nehme E, Weiss LE, Michaeli T, Shechtman Y. 2018. Deep-STORM: super-resolution single-molecule microscopy by deep learning. Optica, OPTICA. 5(4):458–464. doi:10.1364/OPTICA.5.000458.

Nenninger AA, Robinson LS, Hultgren SJ. 2009. Localized and efficient curli nucleation requires the chaperone-like amyloid assembly protein CsgF. Proceedings of the National Academy of Sciences. 106(3):900–905. doi:10.1073/pnas.0812143106.

Olsén A, Arnqvist A, Hammar M, Sukupolvi S, Normark S. 1993. The RpoS Sigma factor relieves H-NS-mediated transcriptional repression of csgA, the subunit gene of fibronectin-binding curli in Escherichia coli. Molecular Microbiology. 7(4):523–536. doi:10.1111/j.1365-2958.1993.tb01143.x.

Olsen WP, Christensen JL, Malle MG, Otzen DE. 2025. Chapter 9 - SMARTQ: single-molecule amyloid fibRil tracking and quantification. A method for accurately imaging, tracking, and quantifying the growth of individual amyloid fibrils using TIRF. In: Tripathi T, Uversky VN, editors. The Three Functional States of Proteins. Academic Press. p. 145–156. https://www.sciencedirect.com/science/article/pii/B9780443218095000065.

Olsen WP, Courtade G, Peña-Díaz S, Nagaraj M, Sønderby TV, Mulder FAA, Malle MG, Otzen DE. 2024. CsgA gatekeeper residues control nucleation but not stability of functional amyloid. Protein Science. 33(10):e5178. doi:10.1002/pro.5178.

Olsen WP, Larsen A-KK, Christensen JL, Malle MG, Otzen DE. 2025. Investigating strategies for creating cross-linked amyloid fibril networks through branching of amyloid growth. Colloids and Surfaces B: Biointerfaces. 251:114617. doi:10.1016/j.colsurfb.2025.114617.

Przybylski A, Thiel B, Keller-Findeisen J, Stock B, Bates M. 2017. Gpufit: An open-source toolkit for GPU-accelerated curve fitting. Sci Rep. 7(1):15722. doi:10.1038/s41598-017-15313-9.

Reinhard S, Ebert V, Schrama J, Sauer M, Kollmannsberger P. 2025. Improving single molecule localisation microscopy reconstruction by extending the temporal context. :2025.04.05.647262. doi:10.1101/2025.04.05.647262. https://www.biorxiv.org/content/10.1101/2025.04.05.647262v1.

Robinson LS, Ashman EM, Hultgren SJ, Chapman MR. 2006. Secretion of curli fibre subunits is mediated by the outer membrane-localized CsgG protein. Molecular Microbiology. 59(3):870–881. doi:10.1111/j.1365-2958.2005.04997.x.

Ruiz-Arias Á, Jurado R, Fueyo-González F, Herranz R, Gálvez N, González-Vera JA, Orte A. 2022. A FRET pair for quantitative and superresolution imaging of amyloid fibril formation. Sensors and Actuators B: Chemical. 350:130882. doi:10.1016/j.snb.2021.130882.

Sage D, Pham T-A, Babcock H, Lukes T, Pengo T, Chao J, Velmurugan R, Herbert A, Agrawal A, Colabrese S, et al. 2019. Super-resolution fight club: assessment of 2D and 3D single-molecule localization microscopy software. Nat Methods. 16(5):387–395. doi:10.1038/s41592-019-0364-4.

Sampson TR, Challis C, Jain N, Moiseyenko A, Ladinsky MS, Shastri GG, Thron T, Needham BD, Horvath I, Debelius JW, et al. 2020. A gut bacterial amyloid promotes α-synuclein aggregation and motor impairment in mice. Chiu IM, Garrett WS, Desjardins M, editors. eLife. 9:e53111. doi:10.7554/eLife.53111.

Sarkar A, Namboodiri V, Kumbhakar M. 2023. Fluorescence correlation spectroscopy measurements on amyloid fibril reveal at least two binding modes for fluorescent sensors. Chemical Physics Impact. 7:100369. doi:10.1016/j.chphi.2023.100369.

Schenk A, Ivanchenko S, Röcker C, Wiedenmann J, Nienhaus GU. 2004. Photodynamics of Red Fluorescent Proteins Studied by Fluorescence Correlation Spectroscopy. Biophysical Journal. 86(1):384–394. doi:10.1016/S0006-3495(04)74114-4.

Shcherbakova DM, Subach OM, Verkhusha VV. 2012. Red Fluorescent Proteins: Advanced Imaging Applications and Future Design. Angew Chem Int Ed Engl. 51(43):10724–10738. doi:10.1002/anie.201200408.

Sofroniew N, Lambert T, Bokota G, Nunez-Iglesias J, Sobolewski P, Sweet A, Gaifas L, Evans K, Burt A, Doncila Pop D, et al. 2025. napari: a multi-dimensional image viewer for Python. doi:10.5281/zenodo.15029515. https://zenodo.org/records/15029515.

Spehar K, Ding T, Sun Y, Kedia N, Lu J, Nahass GR, Lew MD, Bieschke J. 2018. Super-resolution Imaging of Amyloid Structures over Extended Times by Using Transient Binding of Single Thioflavin T Molecules. ChemBioChem. 19(18):1944–1948. doi:10.1002/cbic.201800352.

Speiser A, Müller L-R, Hoess P, Matti U, Obara CJ, Legant WR, Kreshuk A, Macke JH, Ries J, Turaga SC. 2021. Deep learning enables fast and dense single-molecule localization with high accuracy. Nat Methods. 18(9):1082–1090. doi:10.1038/s41592-021-01236-x.

Sun B, Zhou W, Ding T, Porter TS, Wu T, Lew MD. 2023. Single-molecule orientation-localization microscopy resolves how amyloidophilic dye orientations are correlated with amyloid fiber growth and disruption. Biophysical Journal. 122(3):276a. doi:10.1016/j.bpj.2022.11.1575.

Swasthi HM, Basalla JL, Dudley CE, Vecchiarelli AG, Chapman MR. 2023. Cell surface-localized CsgF condensate is a gatekeeper in bacterial curli subunit secretion. Nat Commun. 14(1):2392. doi:10.1038/s41467-023-38089-1.

Thompson RE, Larson DR, Webb WW. 2002. Precise Nanometer Localization Analysis for Individual Fluorescent Probes. Biophysical Journal. 82(5):2775–2783. doi:10.1016/S0006-3495(02)75618-X.

Tuson HH, Biteen JS. 2015. Unveiling the Inner Workings of Live Bacteria Using Super-Resolution Microscopy. Anal Chem. 87(1):42–63. doi:10.1021/ac5041346.

Van Gerven N, Van der Verren SE, Reiter DM, Remaut H. 2018. The Role of Functional Amyloids in Bacterial Virulence. Journal of Molecular Biology. 430(20):3657–3684. doi:10.1016/j.jmb.2018.07.010.

Zhao W, Zhao S, Han Z, Ding Xiangyan, Hu G, Qu L, Huang Y, Wang X, Mao H, Jiu Y, et al. 2023. Enhanced detection of fluorescence fluctuations for high-throughput super-resolution imaging. Nat Photon. 17(9):806–813. doi:10.1038/s41566-023-01234-9.

